# Integrated Cytotoxic and Safety Mechanism of IMV-M™, a MUC16×DR5 Bispecific Antibody

**DOI:** 10.64898/2026.02.10.705083

**Authors:** Victor S. Goldmacher, Iosif M. Gershteyn

## Abstract

**Background:** IMV-M is a MUC16×DR5 bispecific antibody has demonstrated MUC16-selective anti-tumor activity. However, it remained unclear whether multiple binding of IMV-M on a single MUC16 molecule was required for IMV-M cytotoxicity, whether circulating CA125 could attenuate its efficacy or cause off-target toxicity, and whether anti-drug antibodies might induce IMV-M aggregation and related adverse effects.

**Methods:** A comparative analysis of three bispecific antibodies, IMV-M (sofituzumab×DR5), 11D10×DR5, and fluor×DR5, sharing an identical IgG1–anti-DR5 scFv architecture, was performed. Sofituzumab binds to multiple epitopes on a single MUC16 molecule, whereas 11D10 binds a single MUC16 epitope, and fluor does not bind any human antigen. Antibody binding to shed and cell-surface MUC16 was evaluated by ELISA and flow cytometry. Cytotoxicity was assessed in a MUC16^+^/DR5^+^ tumor cell line and MUC16^−^/DR5^+^ hepatic cell lines. Additional studies examined the effects of soluble CA125 and Fc-directed polyclonal antibodies on IMV-M activity.

**Results:** IMV-M bound MUC16 to a markedly higher extent than the 11D10×DR5 comparator, consistent with its multivalent engagement, while binding of fluor×DR5 to MUC16 was negligent. Only IMV-M induced potent cytotoxicity in MUC16^+^ tumor cells, whereas 11D10×DR5 and fluor×DR5 control antibodies were inactive, demonstrating that multivalent clustering on MUC16 is required for apoptosis. IMV-M showed no significant cytotoxicity toward hepatic cell lines, even in the presence of Fc-directed polyclonal antibodies or clinically relevant concentrations of soluble CA125.

**Conclusions:** These findings indicate that IMV-M cytotoxic activity requires clustering on MUC16, that CA125 at clinically relevant concentrations does not mediate IMV-M neutralization, and that aggregate formation with secondary antibodies or soluble MUC16 does not induce off-target toxicity.

## Introduction

Bispecific Apoptosis Triggers (BATs) are a class of bispecific antibodies engineered to simultaneously engage a tumor-associated antigen (TAA) and a death receptor on cancer cells, thereby directly activating the extrinsic apoptotic pathway to induce tumor cell death [1]. By design, BATs address several limitations of current cancer therapies: (i) they selectively target TAAs; (ii) they require neither toxic payloads nor internalization for activity, potentially avoiding the two major limitations of ADCs; (iii) they trigger apoptosis through a mechanism distinct from both ADCs and chemotherapeutics, helping overcome resistance to prior cytotoxic or ADC-based treatments, and (iv) they do not recruit immune cells, allowing BATs to retain activity in the immunosuppressive tumor microenvironment while avoiding the risk of cytokine release syndrome that complicates the use of T-cell engagers and CAR-T therapies.

Initial efforts to induce tumor-selective apoptosis focused on developing BATs that bind both a TAA and DR5. The expectation was that TAA binding would crowd BAT molecules on the cell surface, enabling their DR5-binding domains to drive DR5 clustering and initiate apoptosis. Although several BATs achieved tumor-selective DR5 engagement, early agents underperformed in clinical studies, which could be attributed to insufficient DR5 clustering, resulting in inadequate apoptotic signaling [1]. A significant advance came with the development of IMV-M, a MUC16×DR5 bispecific antibody engineered to cluster DR5 through a unique mechanism that markedly enhances apoptotic signaling [2]. IMV-M was designed to enable clustering of multiple IMV-M molecules on a single MUC16 molecule, resulting in potent DR5 clustering and apoptotic activation. IMV-M has demonstrated robust and selective anti-tumor activity in diverse xenograft models, does not require FcγRIIB-mediated crosslinking (a requirement typical for anti-DR5 IgG antibodies [reviewed in [1]), shows MUC16-selectivity: no activity against MUC16-negative xenografts (a proxy for normal tissues), and displayed no toxicity in the pilot study with two cynomolgus monkeys [2]. However, several key questions remained unresolved: a) although IMV-M’s design implies that clustering on MUC16 is involved in its cytotoxic activity, this had not been directly demonstrated; b) the blood of the majority of patients with ovarian cancer, and some patients with non-small cell lung cancer (NSCLC), or pancreatic ductal adenocarcinoma (PDAC) contains elevated levels of CA125 (shed soluble MUC16) [3-5], which, in principle, could either neutralize IMV-M activity via complex formation, or become toxic to DR5-positive tissues such as liver; c) although IMV-M consists of a humanized IgG1 fused to an scFv derived from a fully human antibody, patients might generate anti-IMV-M antibodies, and such polyclonal antisera could theoretically induce IMV-M aggregation and off-target toxicity. In this manuscript, we extend the characterization of IMV-M by examining whether its cytotoxic activity indeed requires clustering on MUC16, whether shed MUC16, at concentrations typical in the blood of patients, affects IMV-M’s MUC16-dependent cytotoxic activity or induces off-target toxicity, and whether a polyclonal anti-human IgG Fc antibody can induce non-targeted cytotoxicity by crosslinking IMV-M.

## Materials and methods

### Cell lines

NIH:OVCAR-3, HepG2 and Hep3B were obtained from the American Type Culture Collection (ATCC). PK-59 was obtained from Cobioer Biosciences Co.

### Bispecific antibodies

The amino acid sequences of IMV-M and the control proteins are available upon request. The bispecific antibodies for this study were generated at WuXi Biologics using standard transient expression procedures in CHO cells, followed by isolation through protein A chromatography followed by MSS-Superdex 200 chromatography. The integrity and purity of the antibodies were assessed using reduced and non-reduced SDS-PAGE and SEC-HPLC.

### In vitro cytotoxicity testing

The CellTiter-Glo tests (Promega Corporation) were performed at Pharmaron. Cells were plated into flat-bottom tissue culture 96-well plates. The following day, test reagents were added for an additional two days. The relative number of viable cells was then assessed using the CellTiter-Glo assay, following the standard manufacturer’s protocol. All conditions in the in vitro cytotoxicity and apoptosis assays were tested in triplicate, and the mean values ± SD of a representative experiment were plotted. Purified human CA125 (Abbexa, abx060960) contains sodium azide as a preservative. For cytotoxicity assays, sodium azide was removed by desalting CA125 using ThermoFisher 0.5 mL desalting spin columns (89882) pre-equilibrated with culture medium.

### Flow Cytometry Assay

PK-59 cells were used to assess binding of test antibodies via detection of human IgG Fc. The following antibodies were tested: IMV-M, 11D10×DR5, and Fluor×DR5. FITC F(ab’)□ goat anti-human IgG Fcγ (BioLegend, Cat. No. 398006) was used as the secondary antibody. The negative control consisted of cells stained with secondary antibody only. Cell pellets were resuspended at 2 × 10^5^ cells/sample in 95 µL of cold FACS buffer (PBS supplemented with 2% BSA and 2 mM EDTA), followed by addition of 5 µL of 40% goat serum. Test antibodies were prepared in cold FACS buffer at 2× the desired final concentrations. 100 µL of each antibody dilution was added to the corresponding cell suspension (final antibody concentration: 40 nM top concentration, 4-fold serial dilutions, 7 concentrations), and samples were incubated on ice for 1 h. Cells were washed twice with pre-chilled FACS buffer and resuspended in 98 µL of cold FACS buffer. 2 µL of FITC F(ab’)□ goat anti-human IgG Fcγ was added to each sample, mixed thoroughly, and incubated on ice for 30 min protected from light. Cells were then washed once with cold FACS buffer and fixed using pre-chilled 4% paraformaldehyde for 10–15 min on ice. After fixation, cells were washed once with FACS buffer and resuspended in 500 µL of FACS buffer. Samples were analyzed on an iQue3 flow cytometer. Data were analyzed in FlowJo using mean log fluorescence intensity values.

### ELISA (IMV-M, 11D10/DR5, fluor/DR5 binding to CA125)

Human purified CA125 (Abbexa, abx060960) was diluted according to the manufacturer’s instructions and coated onto 96-well plates (100 µL/well) for 1 h at room temperature with shaking (500 rpm). Plates were blocked with a BSA-containing blocking buffer and washed 3× with TBST. Serial dilutions of IMV-M, 11D10/DR5, or the negative control fluor/DR5 were added and incubated for 1 h at room temperature with shaking (500 rpm). After washing, bound antibodies were detected with goat anti-human IgG (γ chain) HRP-conjugated secondary antibody (Invitrogen, 62-8420; cross-adsorbed/depleted against non-human IgG) for 1 h at room temperature with shaking (500 rpm). Plates were washed and developed with TMB (100 µL/well) for 20 min at room temperature with shaking (500 rpm). Reactions were stopped with stop solution, and absorbance was read at 450 nm.

## Results

### IMV-M cytotoxicity depends on multisite engagement of a single MUC16 molecule

To determine whether IMV-M requires simultaneous engagement of multiple epitopes within a single MUC16 molecule to exert cytotoxic activity, we compared two bispecific MUC16×DR5 antibodies, IMV-M (sofituzumab×DR5) and 11D10×DR5, together with a control antibody, fluor×DR5. Sofituzumab binds multiple highly homologous regions (“repeats”) within the extracellular portion of MUC16, whereas 11D10 binds, with affinity similar to sofituzumab, to a single extracellular MUC16 epitope [6]. Fluor×DR5 incorporates an anti-fluorescein IgG that does not bind to human cells [7]. Fluor×DR5 incorporates an anti-fluorescein antibody [7] that does not bind to human cells. All three constructs share an identical architecture: a humanized IgG1 (specific for MUC16 or fluorescein) fused at the heavy-chain C-terminus to the same low-affinity anti-DR5 scFv derived from Lexatumumab (HGS-ETR2) [8] via a flexible linker (Fig. 1).

**Figure 1.**
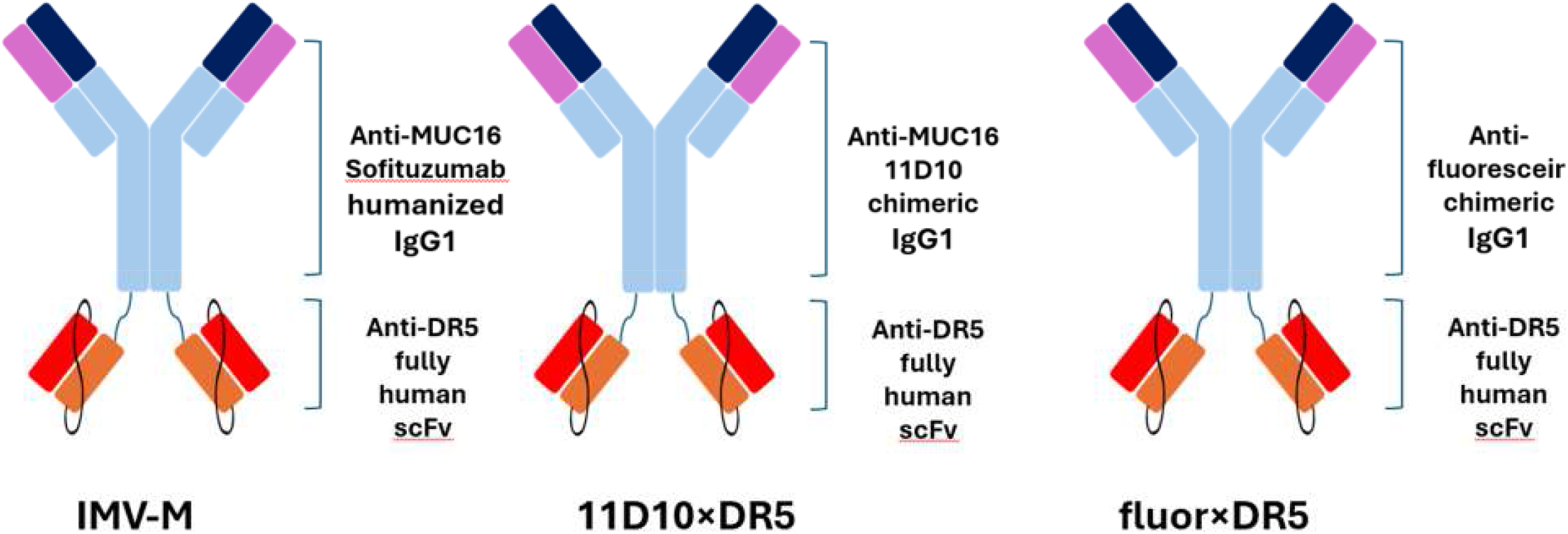
Architecture of IMV-M, 11D10×DR5, and fluor×DR5. Each construct consists of an IgG1 antibody (humanized anti-MUC16 sofituzumab, chimeric anti-MUC16 11D10, or chimeric non-targeting anti-fluorescein IgG) fused at the heavy-chain C-terminus to a low-affinity scFv derived from the anti-DR5 antibody Lexatumumab, connected via a flexible linker.

Two complementary methods were used to compare binding of IMV-M and 11D10×DR5 to MUC16: (i) an ELISA using shed MUC16 (CA125) as the capture reagent, and (ii) flow cytometry assessing binding to MUC16-positive cells detected with the same fluorescent anti-human Fc secondary antibody. IMV-M exhibited ∼11-fold higher binding at saturation to shed MUC16 by ELISA and ∼8.4-fold higher binding to cell-surface MUC16 than 11D10×DR5, consistent with multisite engagement by IMV-M (Fig. 2). Across the concentration range tested, the IMV-M:11D10×DR5 binding ratio increased from ∼2-fold to ∼4-fold and, beginning at ∼2.5×10^−9^ M, reached ∼8-fold (Table 1). Despite these differences in maximal binding, the apparent affinities of the two antibodies were similar in each assay. As expected, fluor×DR5 showed negligible binding to shed MUC16 in ELISA. Fluor×DR5 did bind to cells, but with lower apparent affinity and to a lesser extent than either IMV-M or 11D10×DR5, consistent with exclusive DR5-mediated binding and with the lower affinity of the anti-DR5 scFv for DR5 (0.2 µM [2]) relative to the sub-nanomolar affinities of sofituzumab and 11D10 for MUC16 [6,9].

**Table 1.** Relative binding of IMV-M, 11D10×DR5, and fluor×DR5 to PK-59 cells measured by flow cytometry.

| Concentration, M | IMV-M, MFI<br>minus<br>background* | 11D10×DR5, MFI<br>minus<br>background* | Fluor×DR5, MFI<br>minus<br>background** | MFI<sub>IMV-M</sub>/MFI<sub>11D10×DR5</sub> |
| --- | --- | --- | --- | --- |
| 1 x 10<sup>-11</sup> | 295 | 156 | 0 | 1.9 |
| 4 x 10<sup>-11</sup> | 1207 | 676 | 0 | 1.8 |
| 1.6 x 10<sup>-10</sup> | 3522 | 1639 | 166 | 2.1 |
| 6.3 x 10<sup>-10</sup> | 10021 | 2622 | 267 | 3.8 |
| 2.5 x 10<sup>-9</sup> | 20851 | 2669 | 581 | 7.8 |
| 1 x 10<sup>-8</sup> | 25039 | 3011 | 747 | 8.3 |
| 4 x 10<sup>-8</sup> | 28441 | 3402 | 1021 | 8.4 |

\* Background: MFI of fluor×DR5
\*\*Background: MFI of secondary antibody only

**Figure 2.**
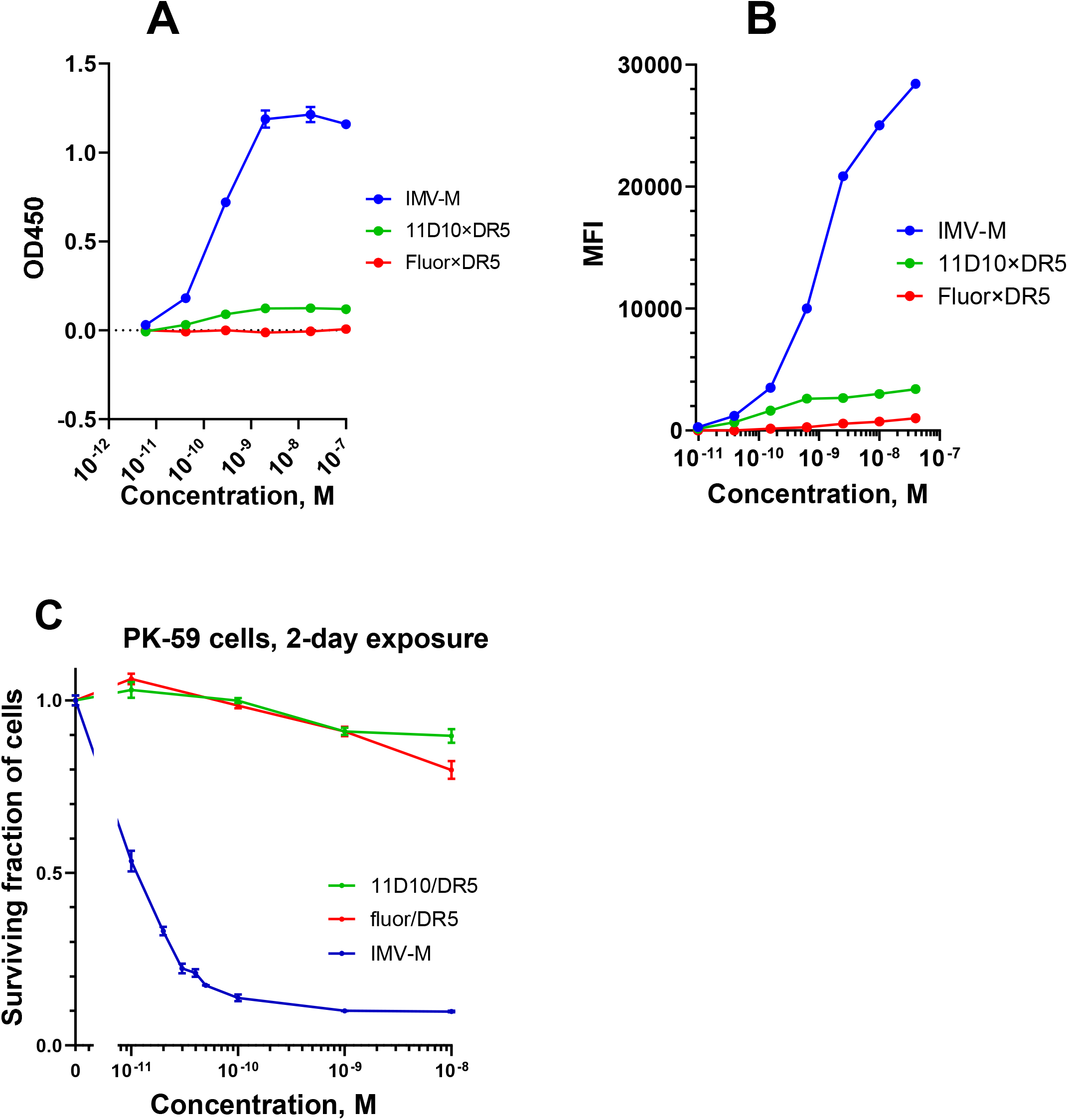
Multisite engagement of a single MUC16 molecule is essential for IMV-M–mediated cytotoxicity. (A) Binding of IMV-M, 11D10×DR5, and fluor×DR5 to shed MUC16 (CA125) purified from human ovarian carcinoma cell lines, measured by ELISA. Data are plotted as OD□□□ (mean ± SD, n= 3) versus antibody concentration; background from secondary antibody–only control was subtracted. (B) Binding to MUC16^+^/DR5^+^ PK-59 cells measured by flow cytometry and shown as mean fluorescence intensity (MFI) versus antibody concentration. (C) Cytotoxicity in PK-59 cells following 2-day treatment, assessed by CellTiter-Glo. Data are presented as mean ± SD (n = 3).

We next evaluated cytotoxicity of IMV-M, 11D10×DR5, and fluor×DR5 in the MUC16^+^/DR5^+^ [2, https://depmap.org/portal] pancreatic adenocarcinoma cell line PK-59. Only IMV-M induced PK-59 cell death (Fig. 2C), with cytotoxicity observed at concentrations as low as 1×10^−11^ M. In contrast, 11D10×DR5 failed to kill PK-59 cells even at concentrations where its cell-surface binding not only matched but markedly exceeded that of IMV-M at 1×10^−11^ M (Table 1). Fluor×DR5 was non-toxic across the entire concentration range tested, indicating that DR5 engagement alone, in the absence of concurrent binding to MUC16, is insufficient to trigger apoptosis in these cells. Together, these results demonstrate that multisite engagement of MUC16, enabling productive clustering, is essential for IMV-M–mediated tumor-cell killing.

### Secondary antibody crosslinking does not elicit IMV-M cytotoxicity in hepatic cell lines

Crosslinking with a secondary antibody has been reported to enhance DR5-mediated cytotoxicity of DR5-targeting agents, presumably by promoting receptor clustering (reviewed in [10]). IMV-M has low affinity for DR5 and is inactive in MUC16-negative xenograft tumors due to the absence of MUC16-mediated clustering [2]. Consistent with this mechanism, fluor×DR5, composed of a non-targeting IgG fused to the same anti-DR5 scFv as IMV-M, also lacks in vivo antitumor activity [2]. Although IMV-M is a humanized IgG1 fused to an scFv derived from a fully human antibody, anti-drug antibodies could potentially develop in patients and, in principle, promote IMV-M clustering and off-target toxicity, particularly in the liver. Normal human hepatocytes are MUC16-negative [11; http://gepia.cancer-pku.cn; www.humanproteomemap] and DR5-positive [12–14]. We therefore tested whether a polyclonal anti-human IgG Fc antiserum could enhance IMV-M cytotoxicity in two MUC16^−^/DR5^+^ human hepatic cell lines, HepG2 and Hep3B (https://depmap.org/portal/). The MUC16^+^/DR5^+^ ovarian cancer cell line OVCAR-3 served as a positive control [2] (https://depmap.org/portal/). Adherent cells were treated for 2 days with IMV-M in the absence or presence of increasing concentrations of rabbit anti-human IgG1-Fc antigen-affinity-purified polyclonal secondary antibody, and cytotoxicity was quantified by CellTiter-Glo. As expected, OVCAR-3 cells were highly sensitive to IMV-M (Fig. 3A). Addition of secondary antibody produced a transition from no change in activity, to modest enhancement at intermediate concentrations, and then reduced cytotoxicity at higher concentrations, defining an optimal crosslinking range for potentiation. In contrast, neither HepG2 nor Hep3B cells exhibited cytotoxicity in response to IMV-M, either in the absence (Fig. 3B) or presence (Fig. 3C,D) of secondary antibody across the entire concentration range tested. These findings suggest a low risk of off-target liver toxicity even if anti-IMV-M antibodies arise during treatment. This conclusion is consistent with the favorable safety profiles reported in clinical trials of prior DR5-targeting antibodies, including Lexatumumab, from which the IMV-M anti-DR5 scFv is derived, and BI 905711, a bispecific antibody incorporating the same anti-DR5 scFv as IMV-M [15]. It is further supported by the favorable safety observed for IMV-M in the pilot study with two cynomolgus monkeys [2].

**Figure 3.**
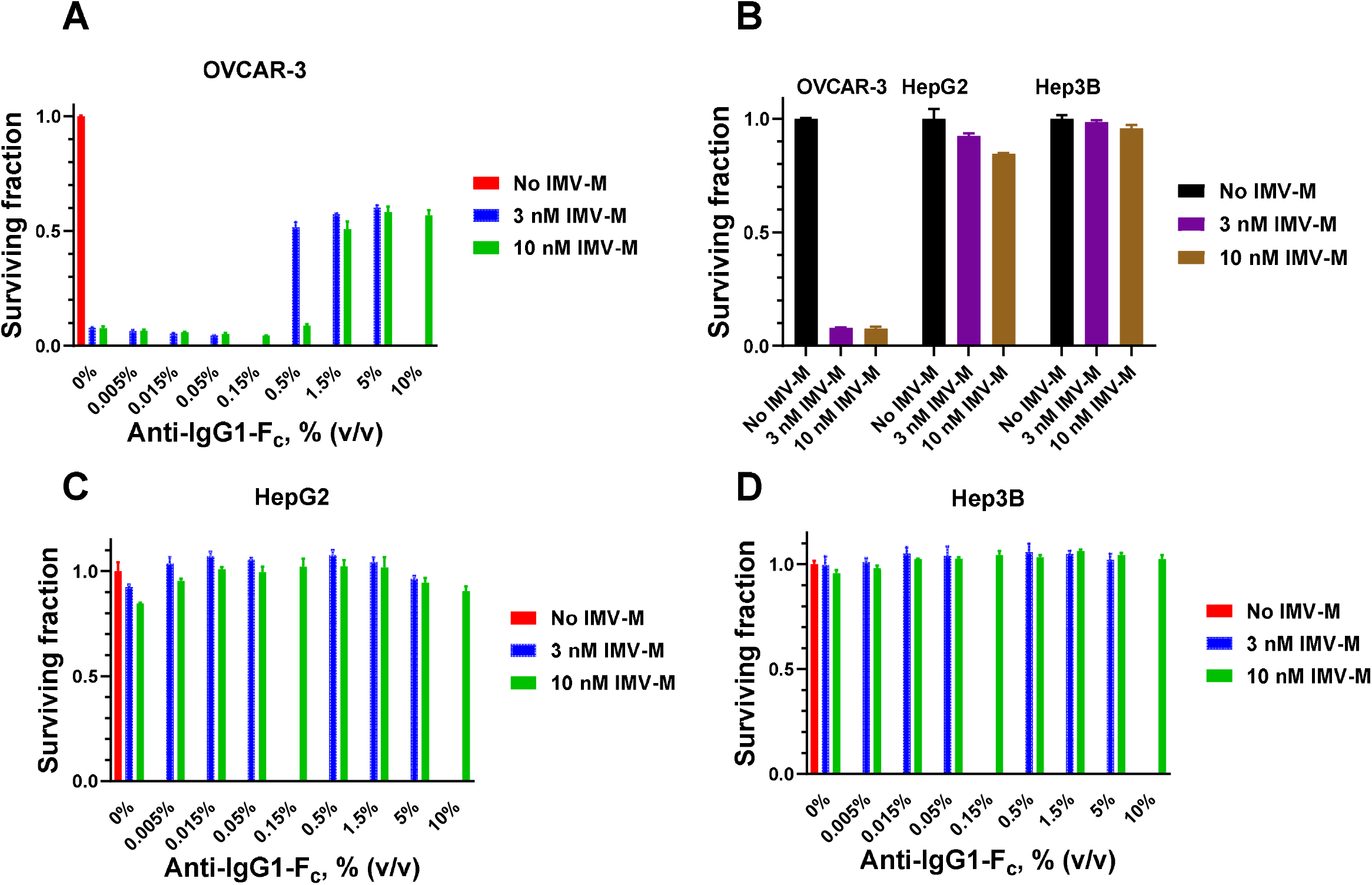
Secondary antibody crosslinking does not confer IMV-M cytotoxicity in hepatic cell lines. (A) Increasing the secondary antiserum:IMV-M ratio modestly enhances IMV-M cytotoxicity toward OVCAR-3 cells at intermediate ratios, whereas higher ratios reduce activity, defining an optimal enhancement range. (B) IMV-M does not induce cytotoxicity in hepatic cell lines. (C, D) IMV-M does not induce cytotoxicity in hepatic cell lines across the entire tested range of secondary antiserum:IMV-M ratios. Cells were treated with IMV-M ± polyclonal rabbit anti-human IgG1 Fc (Sino Biological 10702-T16), and cytotoxicity was assessed by CellTiter-Glo. Data are presented as mean ± SD (n = 3).

### Soluble MUC16 (CA125) neither blocks IMV-M cytotoxicity nor induces significant toxicity in hepatic cells

Patients with ovarian cancer, NSCLC, or PDAC frequently exhibit elevated serum CA125 (shed, soluble MUC16) levels (>35 U/mL). In ovarian cancer, median CA125 levels in blood typically range from ∼70 to 627 U/mL depending on disease stage and histologic subtype and can be substantially higher in some patients [16]. In NSCLC, CA125 is elevated only in a subset of patients and is typically <50 U/mL [17,18]. In PDAC, CA125 levels are often within the normal range but can increase to ∼1,000 U/mL in some patients (and higher in rare cases), depending on disease stage [19,20].

Intravenously administered IMV-M could, in principle, form complexes with circulating CA125, potentially reducing its MUC16-directed activity against MUC16-positive tumor cells and/or generating complexes that might increase hepatotoxicity. To test these possibilities, we evaluated IMV-M cytotoxicity in the MUC16^+^/DR5^+^ pancreatic adenocarcinoma line PK-59 and the MUC16^−^/DR5^+^ hepatic cell line HepG2 in the presence of soluble CA125. Adherent cells were treated for 48 h with IMV-M ± CA125 isolated from conditioned medium of ovarian cancer cell lines, across concentrations representative of those observed in patients. CA125 did not diminish IMV-M cytotoxicity toward PK-59 cells (Fig. 4A), despite detectable binding of IMV-M to this CA125 preparation (Fig. 2A). This suggests that, even at the highest clinically relevant CA125 concentrations tested, the molar abundance of CA125 binding sites is insufficient to sequester IMV-M to a degree that impairs activity. In HepG2 cells, most cells survived IMV-M treatment both in the absence and presence of CA125. Even at the highest CA125 concentration, >50% of HepG2 cells remained viable in the presence of IMV-M. This observation should be interpreted in its physiological context. The adult human liver contains approximately 240 billion hepatocytes [21], whereas 5×10^3^ cells per well were plated in this assay (∼5×10^7^-fold fewer cells). Accordingly, the amount of IMV-M/CA125 complexes per liver cell in 4-6 L of patient’s blood is ∼10^3^-fold lower than the amount of IMV-M/CA125 complexes per cell in a 0.2 mL well in our in vitro experiments. These results suggest that IMV-M is unlikely to induce clinically meaningful liver toxicity via CA125 complex formation.

**Figure 4.**
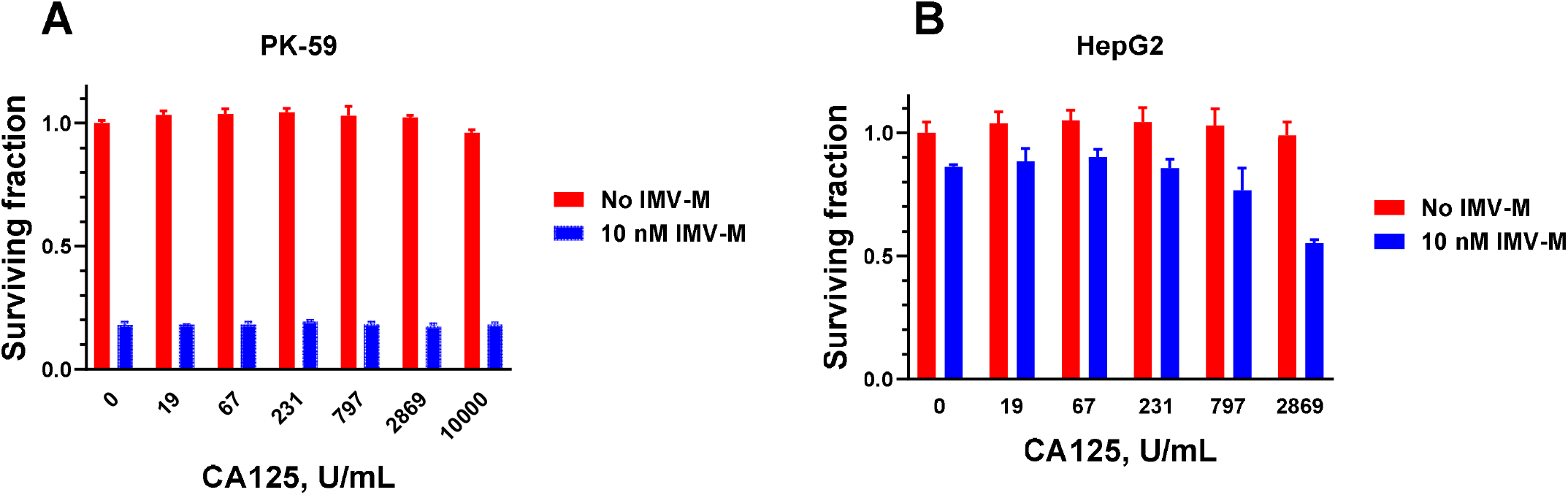
Soluble MUC16 (CA125) does not interfere with IMV-M cytotoxicity and does not confer significant toxicity to HepG2 cells. Adherent PK-59 or HepG2 cells were treated with IMV-M ± increasing concentrations of CA125. Cytotoxicity was quantified by CellTiter-Glo. Data are shown as mean ± SD (n = 3).

## Discussion

One conclusion of this work is that productive DR5 activation by IMV-M requires multisite engagement of MUC16, rather than simple binding to the tumor cell surface. Although IMV-M and 11D10×DR5 share an identical overall architecture and incorporate the same anti-DR5 scFv, only IMV-M induced potent killing of MUC16-positive PK-59 cells. Importantly, 11D10×DR5 remained non-cytotoxic even under conditions where its measured cell-surface binding matched or exceeded that of IMV-M at cytotoxic concentrations. This dissociation between binding and killing argues that overall occupancy alone is insufficient and instead supports a model in which IMV-M must engage multiple epitopes within a single MUC16 molecule to generate the local geometry and avidity needed for effective DR5 clustering. This interpretation is consistent with the markedly higher maximal binding of IMV-M to shed and cell-surface MUC16 relative to 11D10×DR5, despite similar apparent affinities, suggesting that IMV-M achieves increased binding through multisite engagement rather than improved intrinsic affinity. These findings have broader implications for the design of DR5-targeting bispecific antibodies for tumors. DR5 activation is known to require higher-order receptor clustering, and many DR5 agonists have historically shown limited clinical activity, in part due to inadequate clustering in vivo (reviewed in [1,2,10]). Our results suggest that, for bispecific DR5 agonists, mere “crowding” on a tumor-associated antigen (TAA) may not reliably translate into productive DR5 clustering. Instead, an additional mechanism may be required to drive the receptor superclustering necessary for robust apoptotic signaling.

MUC16 undergoes proteolytic cleavage, resulting in shedding of its extracellular domain into the bloodstream. This process generates the cleaved MUC16 fragment known as CA125 [22]. An important question, therefore, is whether circulating shed MUC16 could hinder tumor targeting by IMV-M. Our data indicate that soluble CA125 does not measurably impair IMV-M cytotoxicity toward MUC16-positive tumor cells across a range of CA125 concentrations representative of those observed clinically. Although IMV-M binds shed MUC16 in ELISA, the presence of soluble CA125 at clinically relevant concentrations did not reduce IMV-M–mediated killing of PK-59 cells. These findings suggest that, even in patients with elevated circulating CA125, the abundance of CA125 binding sites is insufficient to sequester IMV-M to an extent that compromises tumor-cell targeting. This is an important translational consideration given the high prevalence of CA125 elevation in ovarian cancer and its elevation in subsets of NSCLC and PDAC patients. This conclusion is further supported by Phase 1 clinical studies of two antibody–drug conjugates (ADCs) that utilize the same anti-MUC16 antibody. Both ADCs demonstrated evidence of clinical activity, including objective tumor responses at doses as low as 0.8 mg/kg and a marked decline in CA125 levels in most patients by day 21 of treatment [23,24]. Together, these observations suggest that circulating shed MUC16 does not neutralize MUC16-directed therapeutics at clinically relevant exposure levels.

Finally, we addressed a clinically relevant safety concern: whether immune-mediated crosslinking or complex formation with soluble antigen could convert IMV-M into an off-target hepatotoxic agent. Hepatotoxicity of TAS266 in some patients, a tetravalent DR5-targeting agent, could be attributed to enhanced cytotoxicity via increasing receptor clustering caused by a pre-existing anti-drug antibodies in these patients [25]. In our study, secondary antibody crosslinking modestly enhanced IMV-M activity in the MUC16-positive control cell line, confirming that the crosslinking conditions were capable of augmenting activity. In contrast, the same crosslinking conditions did not induce cytotoxicity in MUC16-negative hepatic cell lines, consistent with a requirement for MUC16-dependent clustering.

Similarly, CA125 did not confer meaningful cytotoxicity toward HepG2 cells, and quantitative scaling considerations suggest that hepatocyte exposure to IMV-M/CA125 complexes in patients would be substantially lower than in our in vitro assay. These results support the conclusion that IMV-M is unlikely to elicit clinically significant off-target liver toxicity through CA125 complex formation. This interpretation is consistent with the favorable safety observed for IMV-M in the pilot study with two cynomolgus monkeys [2] and with the favorable clinical safety reported for six IgG-based bivalent DR5 antibodies (including one chimeric antibody) and two DR5-bivalent bispecific antibodies [15,26–34], in which immune complex formation may also have occurred in patients.

## Acknowledgements

We thank our colleagues Drs. Yelena Kovtun and Ravi Chari for insightful discussions, and to Dr. Daniel Von Hoff for his critical reading and suggestions regarding the manuscript.

## Statement of Significance

IMV-M requires clustering on MUC16 for an activity. Its potency is not reduced by clinically relevant CA125 levels, and neither CA125 nor secondary antibody induces off-target toxicity, supporting IMV-M’s mechanism of action, safety, and clinical development, and suggesting that antibody crowding on a cell alone may be insufficient for DR5 clustering.

## Data Availability

The data generated in this study are available from the corresponding author upon request.

## Ethics and Consent Statement

consent was not required.

## Animal Research Statement

Not applicable.

